# HiFiTaxa: a reproducible analysis pipeline for full length 16S rRNA gene and ITS amplicon sequencing from PacBio reads

**DOI:** 10.64898/2026.11.03.763332

**Authors:** Alessandro N. Garritano, Flúvio Modolon, Charles Warren

**Affiliations:** School of Life and Environmental Sciences, The University of Sydney, Sydney, NSW 2006, Australia; Centre for Marine Science and Innovation and School of Biological, Earth and Environmental Sciences, The University of New South Wales, Sydney, NSW 2052, Australia; Department of Ecology, Environment and Geoscience, Umeå University, Umeå SE-901 87, Sweden

**Keywords:** 16S rRNA gene, fungal ITS, PacBio HiFi, amplicon sequencing, amplicon sequence variants, GTDB, UNITE

## Abstract

Long-read amplicon sequencing profiles communities at high resolution, but taxonomic accuracy depends on the reference, classifier and the parameters used. Here, we present HiFiTaxa, a reproducible, single command Nextflow pipeline for PacBio HiFi 16S rRNA gene and fungal ITS amplicons. It builds versioned references from current GTDB and UNITE releases and supports Bayesian lowest common ancestor (BLCA) and naive Bayes (NB) classifiers and the read level profilers Emu (16S rRNA gene) and EMITS (ITS). We benchmarked them on mock communities and on holdouts with 10% of GTDB or UNITE sequences withheld. On the 16S rRNA gene mock of 20 species, BLCA detected all 20 species and reported two false positives, NB detected 17 and reported none, and Emu detected all 20 and reported eight. In the GTDB holdout, for taxa represented in the reference, BLCA reached 80.0% species level accuracy against 48.1% for the exact matching species step of NB. However, NB produced fewer incorrect assignments for taxa absent from the reference. On fungal ITS, BLCA had slightly higher holdout accuracy at species level (54.7%) against 52.2% for NB. NB was more conservative for novel taxa, and EMITS detected the most mock species. HiFiTaxa therefore offers a single reproducible workflow that uses current GTDB and UNITE releases and enables users to choose between different classifier algorithms depending on their needs.

## Main text

Long-read sequencing of the full length 16S rRNA gene and internal transcribed spacer (ITS) region^1,2^ resolves lineages that short amplicons cannot^3^. However, accurate taxonomic assignment depends on reference and classifier. The existing workflow has limitations with the PacBio HiFi-16S-workflow github page providing an older Genome Taxonomy Database (GTDB)^4,5^ release (r220), offering only one classifier, without fungal ITS mode and is not maintained anymore. To address these limitations, we developed HiFiTaxa, a single command Nextflow^6^ pipeline that processes from raw HiFi reads to taxonomy (Fig. 1). It rebuilds references from the current GTDB and UNITE^7^ releases. It classifies amplicon sequence variants^8^ with Bayesian lowest common ancestor (BLCA)^9^ and naive Bayes (NB)^10^ and can profile reads directly with Emu^11^ (16S rRNA gene) and EMITS^12^, its ITS counterpart.

**Figure 1.**
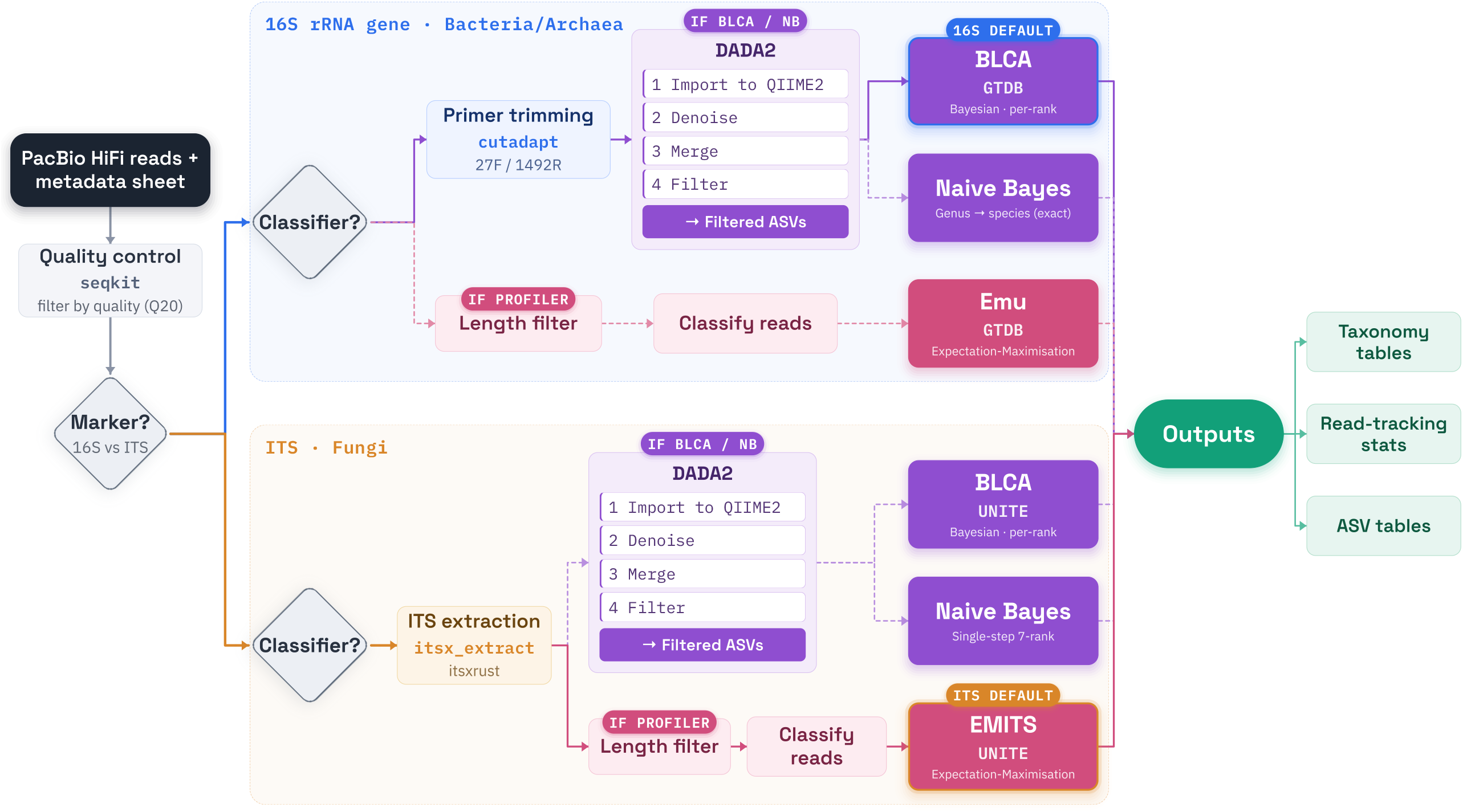
The HiFiTaxa pipeline. A single command Nextflow pipeline that takes raw PacBio HiFi reads and processes to feature counts and taxonomy. For the 16S rRNA gene, primers (27F and 1492R by default; other primer pairs can be supplied) are then removed with cutadapt. For the fungal internal transcribed spacer (ITS), the region is extracted with ITSxRust instead and no primers are trimmed. Classifier marks the choice of classifiers, any combination of which can be run in one command: the BLCA / NB branch infers amplicon sequence variants (ASVs) with DADA2 through QIIME 2 (import, denoising with chimera removal, merging of the per sample tables and the ASV filter) and classifies them, and the IF PROFILER branch length filters the reads and profiles them directly. All classifiers of a marker are anchored on a single reference release: for the 16S rRNA gene the Genome Taxonomy Database small subunit release (GTDB SSU, r232 here) with Bayesian lowest common ancestor (BLCA, the 16S rRNA gene default), the naive Bayes classifier (NB; genus by assignTaxonomy, species by exact matching) and the read level profiler Emu; for ITS, UNITE with BLCA, a single step NB over seven ranks and the read level profiler EMITS (the ITS default). Outputs (right) are a per classifier taxonomy table, a read tracking table and an ASV by sample feature table; Nextflow also writes an execution report and a parameters file for each run.

We validated the 16S rRNA gene path on the ATCC MSA-1003 mock (20 species in 18 genera) in eight Kinnex Revio libraries (Fig. 2a; Supplementary Results and Tables S1 to S5). BLCA and NB detected all 18 genera with no false positive and Emu detected 18 genera, but with four false positives. At species level, BLCA detected all 20 with two false positives, NB detected 17 species with no false positives, and Emu detected 20 species with eight false positives. Both BLCA false positives reflect GTDB circumscription of close relatives (Supplementary Results). Emu reported *Mycobacterium*, absent from the mock, at 5.0%, and *Porphyromonas* and *Escherichia* at 13.7% and 8.7% against 18% expected.

**Figure 2.**
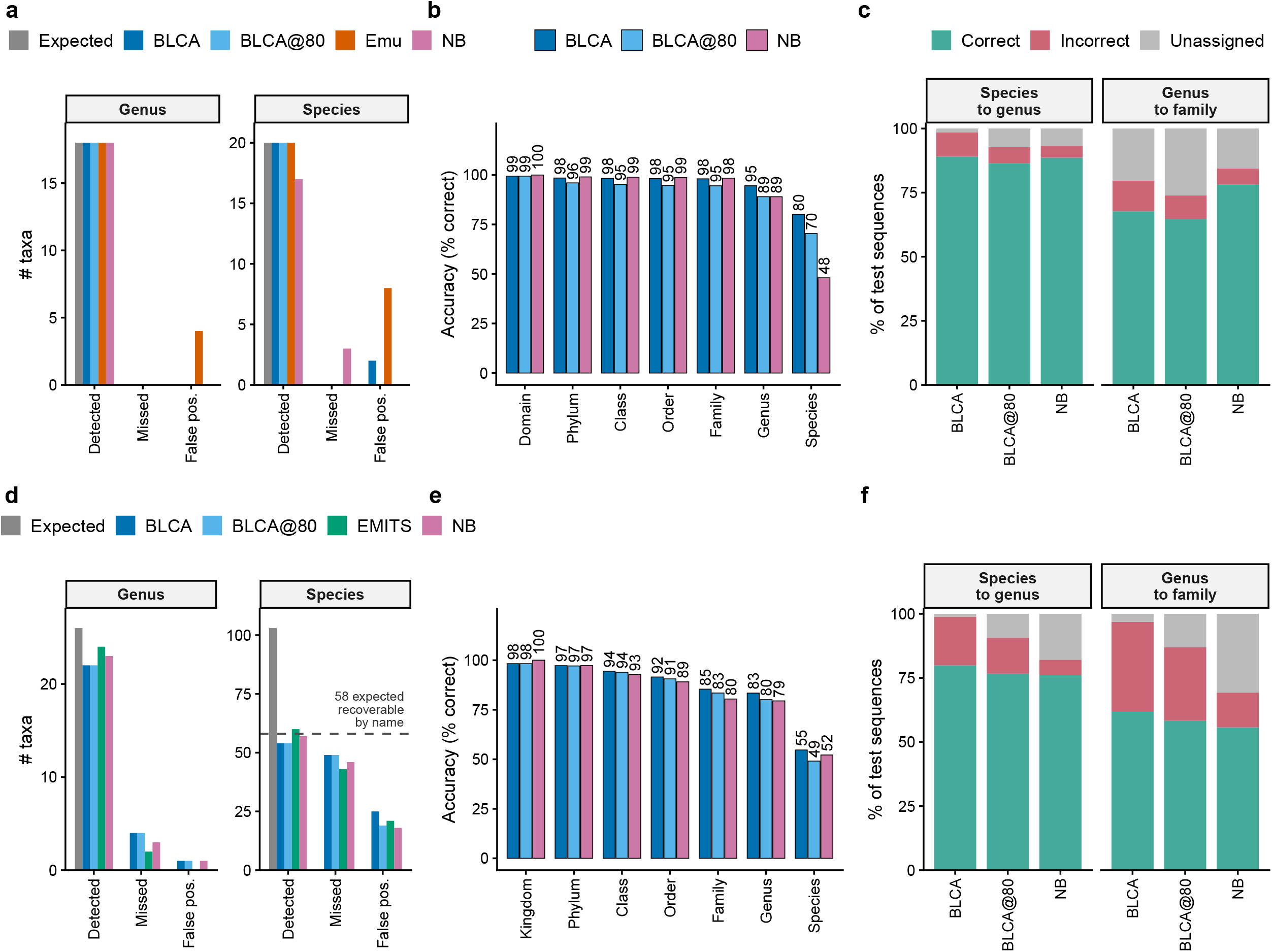
Benchmarking the classifiers on defined mocks and on reference holdouts. Top row (a to c), 16S rRNA gene against GTDB SSU r232; bottom row (d to f), fungal ITS against UNITE. (a, d) Mock community detection. Expected gives the number of reference taxa in the community, and each classifier is scored as Detected (an expected taxon reported at any read abundance above zero), Missed and False positive (a reported taxon carrying more than 0.1% of the reads classified at that rank that is not in the expected composition), at genus and at species. Classifiers are BLCA, BLCA@80 (BLCA with an 80% bootstrap confidence cutoff), the read level profiler (Emu for the 16S rRNA gene, EMITS for ITS) and the naive Bayes classifier (NB). The 16S rRNA gene mock community is ATCC MSA-1003 (20 expected species in 18 genera); the ITS mock is 103 species in 26 genera, and the dashed line in d marks the 58 mock species expected to be recoverable by name from the UNITE release used (Supplementary Methods). Placeholder genera and named synonyms are folded into the corresponding mock member before scoring (Supplementary Methods). (b, e) Per rank holdout accuracy, calculated on the test sequences whose true taxon at that rank (the taxon in their GTDB or UNITE label) is present in the 90% training fraction. For ITS, test sequences with a placeholder label at species or genus rank are excluded at that rank (Supplementary Methods). (c, f) Test sequences whose true taxon is absent from the training fraction, scored at the next rank up: the genus assignment where the species is absent but its genus is present (“species to genus” - left facet), and the family assignment where the genus is absent but its family is present (“genus to family” - right facet). Each assignment is scored as Correct (the correct taxon at that higher rank), Incorrect or Unassigned. For ITS, the placeholder exclusion also applies to these two groups. The read level profilers are absent from b, c, e and f because a holdout contains no reads to profile.

Subsequently, we removed 10% of GTDB r232 (95,731 test sequences - from now on referred as “holdout”) and classified them against a reference database constructed from the remaining 90% of sequences (Fig. 2b, c; Supplementary Tables S6 to S8). Throughout, accuracy is the share of these sequences assigned the correct taxon, with a sequence left without a call counted as a failure, and precision is the share of the calls made that were correct; Supplementary Fig. S1 splits every call into correct, incorrect and unassigned. Both BLCA and NB assigned 98 to 100% correctly from domain to family. At genus level BLCA assigned 94.5% correctly, NB 89.0% and BLCA@80 89.0%; at species level BLCA 80.0%, BLCA@80 70.4% and NB 48.1%. Here, the lower species-level accuracy of NB likely reflects its reliance on exact 16S rRNA gene sequence matches for species assignment.

At genus level precision was 94.8% for BLCA, 96.7% for BLCA@80 and 94.5% for NB. At species level it was 80.3% for BLCA, 90.9% for BLCA@80 and 94.7% for NB, but the three left 0.3%, 22.6% and 49.2% of the sequences without a species call (Supplementary Table S7), so NB was the most precise and the least complete.

Where the closest training sequence was 97 to 99% identical, BLCA assigned the correct genus to 84.7% against 81.7% for NB, and the correct species to 53.2%. Below 97% identity, BLCA assigned the correct genus to 59.0% against 55.2% for NB, and the correct species to 31.2%. NB, which species step requires an identical reference sequence, assigned no species below 100% identity (Supplementary Table S9).

We validated the ITS path on a fungal mock of 103 species in 26 genera^2^ against UNITE (Fig. 2d; Supplementary Methods). BLCA detected 22 of the 26 genera and NB 23, each with one false positive, and EMITS 24 with none. At species level BLCA detected 54 species with 25 false positives, NB 57 with 18 and EMITS 60 with 21. A species counts as detected only when its UNITE name is returned, and for many mock species that could not happen. Of the 103, 7 had no or one PacBio read in the source study and 23 have no entry under their name in the UNITE release used. For another 15 the name is present, but the specimen’s sequence is closer to a UNITE entry labelled with another species than to any entry under its own name, so a classifier returns that other name. That leaves 58 species whose name could be assigned.

In the UNITE holdout (10,213 test sequences), excluding placeholder labels left 1,346 sequences with their species and 6,904 with their genus in the training fraction. Only 7.4% of the sequences scored at species level had a best training hit of at least 99% identity, so this holdout tests new sequences (Supplementary Table S9). BLCA left 1.7% without any hit. However, from class downward BLCA led, with 83.4% at genus level against 79.5% for NB and 54.7% against 52.2% at species level. BLCA@80 gave 80.1% and 49.1%. BLCA assigned an incorrect species to 44.9% and left 0.4% unassigned, whilst NB assigned 15.6% to incorrect species and left 32.2% unassigned. Precision at the genus level was 84.0% for BLCA against 92.8% for NB. At species level, 54.9% for BLCA against 77.0% for NB. Where the species was absent but its genus present, BLCA assigned a wrong genus to 18.9% and NB to 6.0%. On ITS, as on the 16S rRNA gene, BLCA therefore named more sequences correctly at every rank below class and NB made fewer wrong calls.

Broader benchmarks described the trade off between sensitivity and incorrect assignment of taxa absent from the reference^13,14^. In our benchmarks BLCA was the most accurate classifier when the taxon was in the reference and NB made fewer incorrect assignments when it was missing. Incorrect BLCA species assignments concentrated among species with nearly identical 16S rRNA genes (*Staphylococcus aureus* to *S. argentus*).

Many amplicon studies report taxa at genus level. There, BLCA and NB tie on the 16S rRNA gene mock (all 18 genera, no false positive for either), EMITS leads on the fungal ITS mock (24 of 26 genera, no false positive), and the holdouts place BLCA first in accuracy for both markers, with NB the more precise on ITS. HiFiTaxa, therefore provides a pipeline for long-read amplicon analyses in which classifier choice can be guided by evidence rather than by default.

## Supporting information

Supplementary Methods and Results

## Data and code availability

The HiFiTaxa pipeline and its database builders are openly available at https://github.com/alegarritano/HiFiTaxa, and the benchmark job scripts, holdout splitters, scoring and figure code, with the resulting tables and figures, at https://github.com/alegarritano/HiFiTaxa_analysis. The bacterial and archaeal reference databases derive from GTDB SSU r232, publicly available at https://gtdb.ecogenomic.org, and the fungal ITS databases derive from the UNITE general FASTA release for Fungi, version 10.0 of 19 February 2025 (https://doi.org/10.15156/BIO/3301229)^7^. The eight ATCC MSA-1003 PacBio HiFi 16S rRNA gene libraries are from a public Kinnex 16S Revio data set released by PacBio at https://downloads.pacbcloud.com/public/dataset/Kinnex-16S/DATA-ZymoATCCMix-32plex-Revio/, and the 103 species fungal mock reads are those of the source study^2^ (European Nucleotide Archive accession PR-JEB108994).

## Acknowledgements

This research was undertaken with the assistance of resources and services from the National Computational Infrastructure (NCI), which is supported by the Australian Government, under project is54. We thank Torsten Thomas for reviewing the manuscript and for his guidance. The authors acknowledge the use of a generative AI assistant (Anthropic Claude) to assist with pipeline coding, documentation and checking of the analysis scripts; all analytical decisions were made by the authors, who verified the code and take full responsibility for the results.

## Author contributions

A.N.G.: conceptualisation, methodology, software, validation, formal analysis, investigation, data curation, writing (original draft), writing (review and editing), visualisation and project administration. F.M.: formal analysis, writing (review and editing). C.W.: writing (review and editing). All authors approved the final version.

## References

1. Callahan, B. J. et al. High-throughput amplicon sequencing of the full-length 16S rRNA gene with single-nucleotide resolution. Nucleic Acids Research 47, e103 (2019). DOI: 10.1093/nar/gkz569

2. Tedersoo, L. et al. Benchmarking full-length ITS metabarcoding across Illumina 2 × 500, PacBio, and Oxford Nanopore sequencing using mock and soil communities. Molecular Ecology Resources 26, e70189 (2026). DOI: 10.1111/1755-0998.70189

3. Buetas, E. et al. Full-length 16S rRNA gene sequencing by PacBio improves taxonomic resolution in human microbiome samples. BMC Genomics 25, 310 (2024). DOI: 10.1186/s12864-024-10213-5

4. Parks, D. H. et al. A standardized bacterial taxonomy based on genome phylogeny substantially revises the tree of life. Nature Biotechnology 36, 996–1004 (2018). DOI: 10.1038/nbt.4229

5. Parks, D. H. et al. GTDB release 10: A complete and systematic taxonomy for 715 230 bacterial and 17 245 archaeal genomes. Nucleic Acids Research 54, D743–D754 (2026). DOI: 10.1093/nar/gkaf1040

6. Di Tommaso, P. et al. Nextflow enables reproducible computational workflows. Nature Biotechnology 35, 316–319 (2017). DOI: 10.1038/nbt.3820

7. Abarenkov, K. et al. The UNITE database for molecular identification and taxonomic communication of fungi and other eukaryotes: Sequences, taxa and classifications reconsidered. Nucleic Acids Research 52, D791–D797 (2024). DOI: 10.1093/nar/gkad1039

8. Callahan, B. J. et al. DADA2: High-resolution sample inference from Illumina amplicon data. Nature Methods 13, 581–583 (2016). DOI: 10.1038/nmeth.3869

9. Gao, X., Lin, H., Revanna, K. & Dong, Q. A Bayesian taxonomic classification method for 16S rRNA gene sequences with improved species-level accuracy. BMC Bioinformatics 18, 247 (2017). DOI: 10.1186/s12859-017-1670-4

10. Wang, Q., Garrity, G. M., Tiedje, J. M. & Cole, J. R. Naïve Bayesian classifier for rapid assignment of rRNA sequences into the new bacterial taxonomy. Applied and Environmental Microbiology 73, 5261–5267 (2007). DOI: 10.1128/aem.00062-07

11. Curry, K. D. et al. Emu: Species-level microbial community profiling of full-length 16S rRNA Oxford Nanopore sequencing data. Nature Methods 19, 845–853 (2022). DOI: 10.1038/s41592-022-01520-4

12. O’Brien, A., Lagos, C., Fernández, K., Ojeda, B. & Parada, P. EMITS: Expectation-maximization abundance estimation for fungal ITS communities from long-read sequencing. Methods in Ecology and Evolution (2026). DOI: 10.1111/2041-210X.70418

13. Edgar, R. C. Accuracy of taxonomy prediction for 16S rRNA and fungal ITS sequences. PeerJ 6, e4652 (2018). DOI: 10.7717/peerj.4652

14. Bokulich, N. A. et al. Optimizing taxonomic classification of marker-gene amplicon sequences with QIIME 2’s q2-feature-classifier plugin. Microbiome 6, 90 (2018). DOI: 10.1186/s40168-018-0470-z

15. Straub, D. et al. Interpretations of environmental microbial community studies are biased by the selected 16S rRNA (gene) amplicon sequencing pipeline. Frontiers in Microbiology 11, 550420 (2020). DOI: 10.3389/fmicb.2020.550420

16. Turco, S., Giubilei, I., Mahawar, L., Mazzaglia, A. & Albrectsen, B. R. Comparative metabarcoding of ITS1, ITS2, and full-length ITS reveals marker- and tissue-specific variation in fungal community profiling in potato. Plant-Environment Interactions 7, e70168 (2026). DOI: 10.1002/pei3.70168

