## Supplementary Methods and Results for "HiFiTaxa: a reproducible analysis pipeline for full length 16S rRNA gene and ITS amplicon sequencing from PacBio reads"

#### Pipeline implementation

HiFiTaxa is implemented in Nextflow (DSL2)[1] with profiles for Singularity[2] (also used with Apptainer), Docker, conda and a local executor, and the Nextflow version is pinned for reproducibility (24.04 or later). Dependencies run inside containers or conda environments, depending on the profile and the tool. The quality filtering, primer removal and DADA2 denoising modules were adapted from the PacBio HiFi-16S-workflow. The pipeline used QIIME 2 amplicon 2024.10.1, which carries DADA2 1.30.0 and cutadapt 4.9, Emu 3.6.2, ITSxRust 0.2.2 and EMITS 0.1.0; the environments of BLAST+ and minimap2 require versions 2.14 and 2.24 or later. Fig. 1 shows the overall workflow, which branches by marker. Reads from each sample are quality filtered with seqkit[3], which keeps a read when its mean per base error probability over the whole read is at most 1%, reported by seqkit as an average quality of at least Q20. On the 16S rRNA gene path, primers are removed with cutadapt[4] as a linked adapter that requires both primers (maximum error rate 0.1, with the reverse complement of each read also tested so that reads in either orientation are kept), and only reads in which both are found are kept.

#### Primer trimming

By default, HiFiTaxa uses primer pairs 27F- AGRGTTYGATYMTGGCTCAG and 1492R-AAGTCGTAACAAGGTARCY; other primer pairs can be supplied using the flags --forward_primer and --reverse_primer). On the fungal path, the ITS region is extracted from each read with ITSxRust[5] in place of primer trimming; the extraction removes the flanking small subunit and large subunit sequences, which carry the primers.

#### ASV inference

Amplicon sequence variants (ASVs) are then inferred with the DADA2[6] denoise-ccs workflow through QIIME 2[7], with a length window of 1,000 to 1,600 bases applied during 16S rRNA gene denoising and on the Emu[8] read path, and a window of 200 to 1,300 bases for the length variable ITS path. Denoising uses at most two expected errors per read, no minimum base quality, truncation of each read at its first base of quality 2 or lower (truncQ 2, the QIIME 2 default; a truncated read shorter than the length window is dropped), pseudo pooling across samples, one million reads to learn the error model, an OMEGA_C of $10^{-40}$ and the default consensus method for chimera removal; no primer sequences are passed to DADA2, because the primers have been removed or the ITS region extracted beforehand. We chose ASV inference over operational taxonomic unit clustering[9] so that the pipeline resolves sequences at single nucleotide resolution. BLCA and the naive Bayes step classify the DADA2 ASVs, whereas the read level profilers (Emu for the 16S rRNA gene, EMITS[10] for ITS) classify the filtered reads directly without denoising step on DADA2.

#### Reference databases

When the pipeline is launched, HiFiTaxa queries the GTDB release server for the current SSU version and builds the references for all classifiers from a single download of the SSU sequences of all GTDB genomes (the ssu_all file), r232 at the time of writing[11,12], filtered to sequences of at least 1,000 bases. The query is made once at each launch and if a release newer than the local build exists, the launcher asks whether to download and build it. For fungal ITS the pipeline builds from the UNITE general FASTA release for Fungi (version 10.0, 19.02.2025, DOI 10.15156/BIO/3301229)[13], with no minimum length filter. UNITE is not queried and the user downloads the release file from the UNITE repository and the pipeline builds its references from that file. From each source, one build is shared across the classifiers that use it. The BLCA[14] database uses the reference lineage in BLCA format, while the Emu database is built with the build-database command of Emu from the same sequences and a taxonomy map. For the 16S rRNA gene, the naive Bayes references are two files, a genus level (six rank) FASTA for assignTaxonomy and a species assignment FASTA for addSpecies; for ITS a single seven rank FASTA is used. Building the references for all classifiers from one download keeps every classifier on an identical reference.

#### Naive Bayes classification

DADA2[6] is used here not only for denoising but also for the assignTaxonomy function, its implementation of the naive Bayes classifier of Wang et al.[15], and addSpecies, which assigns species by exact sequence matching. For the 16S rRNA gene, the naive Bayes option runs these two functions in a two step process. We ran assignTaxonomy against the genus level reference with minBoot set to 80 and outputBootstraps enabled, which returned assignments from domain through genus with per rank bootstrap confidence, then ran addSpecies against the species reference. A species is assigned only when the ASV matches a reference sequence exactly, the matching references belong to a single species and the genus of that species agrees with the genus assigned by assignTaxonomy. The reverse complement option of addSpecies was enabled; it also tests the reverse complement of each ASV, so that the match does not depend on the orientation of the sequence. For fungal ITS the naive Bayes option runs a single step, a seven rank assignTaxonomy over the full lineage, because the exact match addSpecies step is specific to the 16S rRNA gene references.

#### BLCA and the bootstrap cutoff

BLCA[14] searches each query with BLAST against a BLAST database of the reference sequences, retrieving at most 50 reference sequences per query, and keeps only hits with at least 90% identity and 85% query coverage for the 16S rRNA gene (80% and 50% for ITS). The copy of the BLCA script bundled with HiFiTaxa passes the identity threshold to BLAST itself (its perc_identity option), which the published script fixes at 90% whatever threshold is requested, whereas the ITS analyses use the 80% threshold. It then builds a multiple alignment of the query with the hits whose bit score is within 10% of that of the top hit, and resamples the alignment positions with replacement 100 times. The confidence score at each rank is the percentage of resamples in which the hit most similar to the query belongs to the reported taxon. A query with no hit that passes these filters is left unassigned at every rank. The naive Bayes bootstrap[15] instead draws random subsets of the query in 100 replicates and reports how often it returns the same taxon. Finally, we report BLCA in two forms throughout the benchmark. BLCA, the default that keeps every call, and BLCA@80, the same calls filtered at an 80% bootstrap confidence cutoff. The cutoff trades recall for precision, and reporting both forms separates the effect of the cutoff from the effect of the classifier.

#### Outputs

A single run writes one output directory. It holds read tracking tables that follow counts through primer removal (16S rRNA gene only) and through DADA2 filtering, denoising and chimera removal; the inferred ASV sequences with an ASV by sample feature table; and a taxonomy table for each classifier selected. BLCA returns a seven rank taxonomy per ASV with per rank bootstrap confidence, for both markers. The naive Bayes step returns a per ASV genus with bootstrap confidence and, for the 16S rRNA gene, an exact match species, together with the ASV frequency table merged with that taxonomy. Because it is a sequence comparison and not a bootstrap estimate, specie-level assignments carry no bootstrap value. Emu returns a relative abundance table per sample, which the pipeline also combines into genus level and species level relative abundance matrices of samples by taxa; EMITS returns genus level and species level relative abundance tables per sample. The per sample tables give the relative abundance and the estimated number of reads for each taxon. Nextflow also writes an execution report for each run that records the command line, the profile, the Nextflow version, the run time and the resources used by each task; a summary of the main run settings is written to a separate file (parameters.txt).

#### Holdout benchmark

To assess classifier accuracy, we performed a sequence holdout benchmark on GTDB SSU r232. After a 1,000 minimum-length filter, the reference was split at random (seed 42) into a 90% training fraction (861,581 sequences) and a 10% test set (95,731 sequences). We then classified the test sequences and compared each call to its GTDB taxonomic label. Because a holdout contains no reads to profile, only the ASV classifiers (BLCA, BLCA@80 and naive Bayes) were run on the holdout. We report accuracy for the test sequences whose true taxon at the rank in question is present in the training fraction (Supplementary Fig. S1 and Supplementary Table S6).

At species level, these are 90,688 sequences. Accuracy at a rank is the number of test sequences assigned the correct taxon divided by the number of test sequences whose true taxon at that rank is present in the training fraction. Because the reference was split at the sequence level and GTDB holds many identical SSU sequences, a test sequence may have an identical sequence in the training fraction.

To measure this, we first found the test sequences identical to a training sequence by exact comparison (74,673 of the 95,731 16S rRNA gene test sequences, 78.0%; none of the 10,213 ITS test sequences) and then searched the remaining sequences against the training fraction with BLAST (at most five hits per query, minimum identity 80%), keeping the best hit with at least 80% query coverage. Test sequences were classified by the identity of that hit (100%, 99 to under 100%, 97 to under 99%, under 97%, no hit) and the accuracy of each classifier was calculated within each bin on the same subsets as above (Supplementary Table S9).

For test sequences whose taxon was absent from the training fraction (Fig. 2 and Supplementary Table S8), we asked whether each classifier still assigned the correct taxon at the next rank up. We grouped the 5,043 sequences whose species was absent by whether the next higher rank was still present in the training fraction. 4,130 with the species absent but the genus present, 771 with the genus absent but the family present, and the remaining 142 with neither the genus nor the family present, which were not scored. Each group was scored one rank up from the absent taxon. Sequences with the species absent but the genus present were scored at genus, and sequences with the genus absent but the family present at family.

Each group is reported as the fractions correct, incorrect and unassigned, so that assigning the correct taxon at the nearest rank still present in the training fraction counts as correct. The fungal ITS holdout was built the same way from the UNITE reference[13], split (seed 42) into a 91,924 sequence training fraction and a 10,213 sequence test set, and scored in the same way, except for ASVs corresponding to species placeholders (for instance *Tomentella* sp.) or of a genus name (labels containing Incertae sedis).

A match to those labels does not show that the species or genus was identified, so the 6,248 test sequences with a placeholder species label were excluded from scoring at species level and the 3,101 with an Incertae sedis genus label from scoring at genus level. This left 1,346 test sequences whose species was present in the training fraction and 6,904 whose genus was present. Sequences with a placeholder species label were also excluded from the group with the species absent but the genus present, and sequences with an Incertae sedis genus label from both groups scored one rank up, which left 2,420 and 192 sequences in the two groups. The exclusion was not applied to the 16S rRNA gene holdout, because GTDB placeholder names denote species defined by genomes.

#### Classifier agreement on the holdout

For every 16S rRNA gene holdout sequence, we compared the BLCA and naive Bayes genus calls with each other and with the GTDB annotation, and scored the pair as both correct, BLCA correct only, naive Bayes correct only, both wrong on the same genus, both wrong on different genera, or unassigned when at least one classifier made no assignment and neither was correct. Sequences were then grouped by their true genus and split into those whose species is present in the training fraction and those whose species is absent. Supplementary Tables S10 and S11 report every genus carrying at least one test sequence. The corresponding figures (Supplementary Figs. S2 and S3) plot the genera with the lowest agreement between the classifiers, among those with at least five test sequences and at least one sequence on which the two classifiers were not both correct, up to one hundred genera (100 of 305 in Supplementary Fig. S2 and all 28 in Supplementary Fig. S3).

#### Mock community

Eight publicly available PacBio HiFi libraries of the ATCC MSA-1003 staggered mix of 20 strains, which spans four abundance tiers from 18% down to 0.02%, were processed through HiFiTaxa against the GTDB SSU r232 database. These libraries are from a public Kinnex 16S Revio data set released by PacBio (forward barcodes 01 to 08 paired with reverse barcodes 37 to 44, listed as the ATCC staggered mix in the sample key of the data set; `https://downloads.pacbcloud.com/public/dataset/Kinnex-16S/DATA-ZymoATCCMix-32plex-Revio/`), which is distributed as a direct download.

The libraries hold 2.9 to 4.0 million reads each (27,080,353 in total), all of which were processed. The quality filter removed 14.6% of them and read counts after each step are given in Supplementary Table S4. For the mock community, the two ASV filters of the pipeline were disabled (minimum total read count and minimum number of samples both set to 0), so that mock members expected at 0.02% were retained. In real data this setting may lead to false positives, but here it was kept by suggestion of the original PacBio developers to assess the accuracy on the full dataset and compare with the original pipeline performance. HiFiTaxa's defaults are a minimum total of 5 reads across all samples and a minimum of 1 sample.

Observed composition is the relative abundance across the eight libraries, taken for Emu as the mean over all eight libraries of the per sample values in its genus and species tables, with a taxon that Emu did not report in a library counted as 0 for that library, and for BLCA and naive Bayes by summing ASV read counts over all libraries at each rank and dividing by the total read count.

Before scoring, we folded GTDB placeholder genera and named GTDB synonyms of mock members into the corresponding mock organism, both for presentation and for counting false positives. These include the placeholder genera split from *Escherichia coli*, which were folded to *Escherichia*, and the genus *Pauljensenia*, under which GTDB reports the mock member *Schaalia odontolytica* as *Pauljensenia dentiphila*; we folded the genus to *Schaalia* and the species to *Schaalia odontolytica*, since GTDB and the National Center for Biotechnology Information (NCBI) assign that same genome different names. Both *Schaalia* and *Pauljensenia* were proposed in the 2018 genome based reclassification of the *Actinobacteria*[16]. Because these placeholders fold to a genus that is present in the mock, sequences carrying them count as detection of that genus and not as false positives. At species level, a call was credited to a mock member only when it was the name under which ATCC lists that strain, its NCBI name or its GTDB r232 name.

*Escherichia coli* K-12 MG1655 was therefore also credited under its GTDB r232 name, G047199095 sp047199095, *Bacillus cereus* ATCC 10987 under *Bacillus_A pacificus* and *Pseudomonas aeruginosa* ATCC 9027 under *Pseudomonas paraeruginosa. Escherichia boydii* and ECMA0423 sp047199055 are GTDB r232 species that do not contain the *E. coli* strain of the mock, so they were counted as false positives where they exceeded the threshold given below (*E. boydii* for BLCA, eight ASVs with 5.4% of reads together; ECMA0423 sp047199055 for Emu, 1.2%). For the same reason we did not fold *Staphylococcus argentus* into *S. aureus.* Although GTDB circumscribes *S. argenteus* to include genomes deposited at NCBI as *S. aureus*, one of which is present in the reference (GCA_008619595.1), *S. argenteus* is a distinct validly published species, and folding two valid species would overstate specificity. Therefore, we counted the single *S. argentus* call as a false positive.

A taxon is considered detected when a classifier reports it at any read abundance above zero. A false positive is a foreign taxon, one not in the expected composition after folding, reported above 0.1% of the reads classified at that rank. Reported taxa that lack a genus or species name (for example *Cortinarius* sp. in the fungal mock) were not counted as false positives. False positive counts at thresholds of any read and of 0.01% of the classified reads are given in Supplementary Table S5. Emu reported *Mycobacterium mageritense* at 4.5 to 5.4% in every library although no *Mycobacterium* is in the mock. The GTDB r232 genome GCF_020884715.1 (GenBank assembly GCA_020884715.1, a scaffold level assembly with a CheckM contamination estimate of 4.07%) is assigned to *M. mageritense* and is not a species representative. Besides its own 16S rRNA gene, its SSU sequences in the GTDB ssu_all file include one (contig NZ_BPWM01000103.1) identical over 1,528 bases to the 16S rRNA gene of *Porphyromonas gingivalis* and one (contig NZ_BPWM01000098.1) 99.6% identical over 1,217 bases to that of *Escherichia coli*, as found by searching the *M. mageritense* SSU sequences against the GTDB r214 ssu_all file with BLASTn; the *P. gingivalis* copy is also present in r232, where the holdout identity search returned the *P. gingivalis* genome NZ_CP007756.1 as its identical best training hit over 1,528 bases. Both species are mock members at 18%, and Emu reported them below expectation (13.7% and 8.7%), whereas BLCA and NB reported no *Mycobacterium.* We therefore attribute the *M. mageritense* calls to these contaminant sequences and not to the reads.

#### Fungal ITS mock

The 103 species fungal mock (26 genera) was sequenced on PacBio Revio by the source study with the primers ITS9MUNngs and ITS4ngsUni, which amplify the full length ITS region[17,18]. HiFiTaxa does not trim primers on the ITS module. The ITS region is extracted from each read with ITSxRust whatever primers were used. The two libraries hold 19,722 and 32,411 reads (European Nucleotide Archive runs ERR16773545 and ERR16773546), of which 19,458 and 31,822 extracted ITS sequences entered DADA2 (Supplementary Table S4). We processed the mock through the pipeline to ITS amplicon sequence variants, with the two ASV filters at a minimum total of one read in one sample, which retained all 138 ASVs (44,758 reads, the non chimeric total of Supplementary Table S4), and classified it with BLCA and the single step naive Bayes classifier against the full UNITE reference. We also ran EMITS (`https://github.com/ayobi/emits`), an expectation maximisation classifier for ITS analogous to Emu, at the read level.
The ITS sequences extracted from the mock reads were mapped to UNITE with minimap2[19] under its map-hifi preset, and EMITS was run on the alignments under its pacbio-hifi preset.

Reported taxa were folded to mock members as for the 16S rRNA gene mock. The genera *Thaxterogaster*, *Phlegmacium* and *Calonarius*, which a revised classification separated from *Cortinarius*[20], were folded into *Cortinarius.* Two mock members that the classifiers reported are present in the UNITE release only under a variant of their name, *Cortinarius ochraceobrunneus* as *Phlegmacium ochraceobrunneum* and *Calocybe obscurissima* as *Rugosomyces obscurissimus.* Each variant was folded into the mock member, and the genus *Rugosomyces* into *Calocybe.*

The fungal mock was prepared at an equimolar DNA content, so has no abundance ground truth. It is, therefore, assessed by detection rather than by composition. The source study recovered 93 of the 103 mock species from its PacBio reads with its minimum and standard filtering pipelines, and 92 with ASVs[18], against 54 to 60 species detected here. The two counts come from different tests. Tedersoo et al. counted a mock specimen as recovered when a representative sequence had up to two mismatches to the Sanger sequence of that specimen, whereas we counted a species only when a classifier returned its name from UNITE.

To compare the two, we searched the Sanger sequence of each specimen against the UNITE release with BLASTn. Of the 103 mock species, 6 had no PacBio reads in any pipeline of the source study, 1 had a single read, a further 23 have no entry under their name in the UNITE release used, and for another 15 the best BLASTn hit of the specimen sequence in UNITE is a species hypothesis carrying a different label.

The remaining 58 species are those expected to be recoverable by name; of these, BLCA, NB and EMITS named 54, 54 and 58. NB and EMITS also returned the names of three and two mock species outside these 58. NB *Cortinarius majoranae* (reported as *Phlegmacium majo-*

*ranae*), *Leccinum scabrum* and *Russula amethystina*, and EMITS *Cortinarius semisanguineus* and *Russula amethystina*. As mock members, none of the five is counted as a false positive. Most false positive species (19 of 25 for BLCA, 14 of 18 for NB and 19 of 21 for EMITS) carry the label of the UNITE species hypothesis that is the best BLASTn hit of a mock specimen. At genus level there is no such gap, and we detected 22 to 24 of the 26 genera, and the PacBio pipelines of the source study 23 or 24.

##### Comparison with the PacBio pipeline

The NB classifier of HiFiTaxa follows the PacBio HiFi-16S-workflow (`https://github.com/pacificbiosciences/HiFi-16S-workflow`) in read filtering, primer removal, denoising with DADA2 and classification with the assignTaxonomy function at a bootstrap cutoff of 80. It differs in the reference and in the species step, as HiFiTaxa uses one GTDB release, with a genus level training file and exact matching for species, whereas the PacBio workflow runs assignTaxonomy against species level training files of Greengenes2[21], GTDB and SILVA, first at species and then at genus level, and merges the three results.

#### Supplementary Results

The 16S rRNA gene mock, ATCC MSA-1003, comprises 20 species across 18 genera and was profiled in eight libraries (Fig. 2a; Supplementary Fig. S4 and Supplementary Tables S1 and S2). Of the 23.1 million reads that passed the quality filter, 99.9% carried both primers and 47.3% passed the DADA2 filter of at most two expected errors per read and chimera removal, so 10.9 million reads entered the 246 ASVs (Supplementary Table S4). Both BLCA species level false positives were close relatives of mock members. *Staphylococcus argenteus* (0.4% of reads) is a validly published sister species of *S. aureus* that GTDB circumscribes to include a genome deposited at NCBI as *S. aureus*. *Escherichia boydii* (5.4% of reads) is a GTDB species that does not include the *E. coli* strain of the mock (Supplementary Methods). At the 80% bootstrap cutoff BLCA still detected every genus and species, but it left 19.9% of the reads without a genus, including 85% of the Escherichia reads (18.4% without the cutoff against 2.7% with it; Supplementary Table S3). GTDB r232 places genomes formerly labelled E. coli in three genera, Escherichia and the placeholder genera ECMA0423 and G047199095, the last holding the mock strain K-12 MG1655, so the bootstrap splits its support among them.

Emu detected all 18 genera and all 20 species, although it reported *Cutibacterium acnes* at 0.0002% of reads, against 0.18% expected, and in only two of the eight libraries. It also reported eight false positive species and was the only classifier to report false positive genera, four in total. The most abundant of these, *Mycobacterium* (a genus absent from the mock), reached a mean relative abundance of 5.0%, and Emu reported *Escherichia* at 8.7% and *Porphyromonas* at 13.7%, against 18% expected for each. The *Mycobacterium* calls and the *Porphyromonas* shortfall is likely due to the GTDB reference.

GTDB r232 genome GCF_020884715.1, labelled *Mycobacterium mageritense*, carries besides its own 16S rRNA gene an SSU sequence identical over 1,528 bases to the 16S rRNA gene of *Porphyromonas gingivalis* and one 99.6% identical to that of *Escherichia coli* (Supplementary Methods). Emu, which apportions every read among the references it matches, assigned part of the *P. gingivalis* reads to *M. mageritense* in every library (4.5 to 5.4%), and *Porphyromonas* and *Mycobacterium* together came to 17.8 to 19.7% per library (mean 18.7%), close to the 18% expected for *Porphyromonas*. BLCA and NB were unaffected because the many correctly labelled reference sequences of both species outweigh the two contaminant copies. However, for *Escherichia* Emu reported *Escherichia* at 6.5 to 11.1% per library against 14.0 to 22.7% for BLCA, a gap larger than the *Mycobacterium* calls and uncorrelated with them across libraries (Supplementary Table S12). BLCA and NB both reported *Porphyromonas* at 16.3%, and *Escherichia* at 18.4% and 15.8%, respectively. All three classifiers reported *Cereibacter* close to the 18% expected (16.3% for BLCA and NB and 18.6% for Emu; Supplementary Table S3 and Supplementary Fig. S5). Summed over the 18 genera and the genera absent from the mock, the absolute deviation of the observed from the expected composition was 18.7 percentage points for BLCA, 20.5 for NB, 30.0 for BLCA@80 and 36.3 for Emu (Supplementary Table S3). Part of this deviation is shared by every classifier, for example *Bacillus* at 4.5 to 5.5% against 1.8% expected and *Streptococcus* at 14.5 to 15.6% against 19.8%, likely reflecting the amplicon biases rather than the classifiers. The false positive counts in Fig. 2a use a threshold of 0.1% of reads. Counting every foreign genus reported at any read abundance gives 15 for BLCA, 12 for NB and 78 for Emu, and the four Emu genera are the only ones that remain above 0.1% (Supplementary Table S5).

In the 16S rRNA gene holdout (n = 95,731 test sequences; Fig. 2b, Supplementary Fig. S6 and Supplementary Table S6), 90,688 test sequences had their species present in the database. BLCA and NB assigned between 98 and 100% of the test sequences whose taxon was present correctly at each rank from domain to family, and BLCA@80 between 94.5 and 99.4%. At genus level BLCA and NB assigned an incorrect genus at a similar rate (5.2% and 5.1%), and BLCA@80 at a lower rate (3.0%; Supplementary Table S6). BLCA@80 uses the same cutoff of 80% that NB applies through the DADA2 minBoot setting.

With the cutoff, BLCA@80 kept a species call for 77.4% of the test sequences whose species was present in the training fraction, and 90.9% of the calls it kept were correct (its precision; Supplementary Table S7). The precision of BLCA, run without a cutoff, was 94.8% at genus and 80.3% at species. NB's was 94.5% and 94.7%, so the two were equally precise at genus level, whereas at species level NB was the more precise but assigned only half of the sequences (Supplementary Table S7). At species level BLCA assigned an incorrect species to 19.7% of sequences and left almost none unassigned, whereas NB assigned an incorrect species to 2.7% and left 49.2% unassigned (Supplementary Fig. S1). The lower accuracy of NB therefore reflects sequences left unassigned, not incorrect assignments.

On the 16S rRNA gene this is a contrast between BLCA and exact matching, not between

BLCA and the naive Bayes classifier. Exact matching can assign a species only when a sequence that matches the test sequence exactly remains in the training fraction. Because GTDB holds many identical 16S rRNA gene sequences, 74,673 of the 95,731 test sequences (78.0%) have a sequence identical to a training sequence, so for these sequences the holdout tests the retrieval of an identical sequence rather than the classification of a new one. BLCA, which is run without a confidence cutoff, leaves 0.6% of the 16S rRNA gene test sequences and 1.7% of the ITS test sequences without any assignment, and 0.3% and 0.4% without a species (Supplementary Table S6).

The incorrect species assignments of BLCA were concentrated among closely related species with nearly identical 16S rRNA genes, for example *Staphylococcus aureus* assigned to *S. argenteus*. The BLAST search of BLCA may contribute to these errors.

Most 16S rRNA gene test sequences have a near identical training sequence (Supplementary Table S9). Those with species present in the training fraction, 84.3% had a best training hit of 100% identity, 14.1% one of 99 to under 100%, 0.9% one of 97 to under 99%, 0.6% one below 97% and 0.1% no hit. Genus accuracy fell with identity for every classifier, from 95.5% (BLCA), 89.5% (NB) and 89.4% (BLCA@80) at 100% identity to 84.7%, 81.7% and 81.0% at 97 to under 99% and 59.0%, 55.2% and 54.7% below 97%. At species level BLCA assigned 80.1% correctly at 100% identity, 84.3% at 99 to under 100%, 53.2% at 97 to under 99% and 31.2% below 97%; NB assigned a species to 57.1% of the sequences with an identical hit and to none of the others, because its species step requires an exact match.

For the 5,043 test sequences whose species was absent from the training fraction, we asked whether each classifier still assigned the correct taxon at the next rank up (Fig. 2c and Supplementary Table S8). Where the species was absent from the training fraction but its genus was present (4,130 sequences), BLCA and NB assigned the correct genus at similar rates (89.1% and 88.6%), but BLCA had more than double the rate of incorrect genus calls (9.5% against 4.5%). Where the genus was absent but its family was present (771 sequences), NB assigned the correct family for 78.2% and BLCA for 67.7%. NB therefore makes fewer incorrect assignments in both groups, while BLCA assigns the correct genus to a similar share of the sequences whose species is absent, with a higher rate of incorrect assignments.

On the fungal ITS mock of 103 species across 26 genera, EMITS was the most sensitive method, detecting 24 of 26 genera and 60 of 103 species ahead of NB (23, 57) and BLCA (22, 54), with no false positive genus compared with one for each of the others (Fig. 2d). That lower number of false positives did not carry through to species, where EMITS returned 21 false positives compared with 18 for NB and 25 for BLCA, and 20 of those 21 fell in the species rich genera *Cortinarius* and *Russula*. The source study recovered 84 to 96 of the 103 species from its PacBio reads, depending on the pipeline[18], but it scored a specimen by the match of a sequence to its Sanger sequence and not by name.

Of these 58 recoverable by name (see Supplementary Methods), EMITS named 58, NB 54 and

BLCA 54; the five further names that NB and EMITS returned belong to mock species outside the 58 (Supplementary Methods), so the detection counts in Fig. 2d are bounded by the reference and the source data rather than by the classifiers.

For the UNITE holdout, from class downward, BLCA led at every rank, reaching 83.4% at genus level against 79.5% for NB, although at species level the two were close (54.7% for BLCA and 52.2% for NB); with the matched 80% cutoff, BLCA@80 reached 80.1% at genus and 49.1% at species. At species level BLCA also assigned an incorrect species to 44.9% of the test sequences whose species was present in the training fraction and left 0.4% unassigned, whereas NB assigned 15.6% incorrectly and left 32.2% unassigned; at genus level the unassigned fractions were 0.7% for BLCA and 14.4% for NB (Supplementary Table S6). The precision of the calls made was 54.9% for BLCA and 77.0% for NB at species level, and 84.0% against 92.8% at genus level (Supplementary Table S7).

### Supplementary Figures

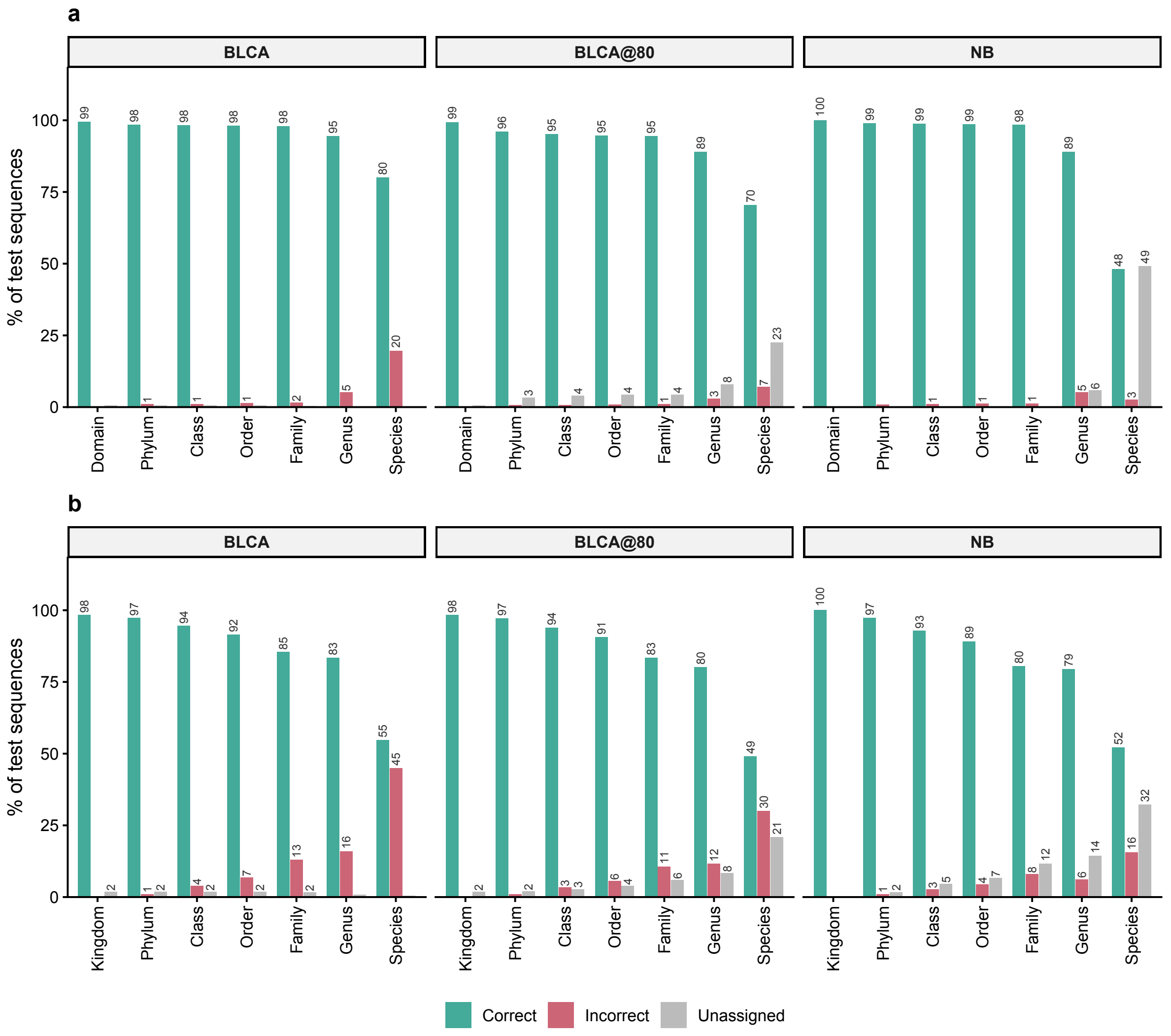


**Supplementary Figure S1. Breakdown of the holdout calls into correct, incorrect and unassigned.** For (a) the 16S rRNA gene GTDB r232 holdout and (b) the fungal ITS UNITE holdout, every call on the test sequences whose true taxon at that rank is present in the 90% training fraction is scored as Correct (the correct taxon; this bar is the per rank accuracy), Incorrect (a wrong taxon was assigned) or Unassigned (no call was made). The three outcomes are drawn as side by side bars at each rank for BLCA, BLCA@80 and the naive Bayes classifier (NB), and sum to 100% at every rank. Values below 1% are left unlabelled. For ITS, test sequences with a placeholder label at species or genus rank are excluded at that rank (Supplementary Methods). The breakdown separates a classifier that misclassifies from one that leaves sequences unassigned. At species level BLCA is 80.0% correct but 19.7% incorrect with almost nothing unassigned, whereas NB is only 2.7% incorrect and leaves 49.2% unassigned, because its species assignments on the 16S rRNA gene require an exact match to a reference sequence. The read level profilers Emu and EMITS are absent as a holdout contains no reads to profile. Underlying values are in Supplementary Table S6.

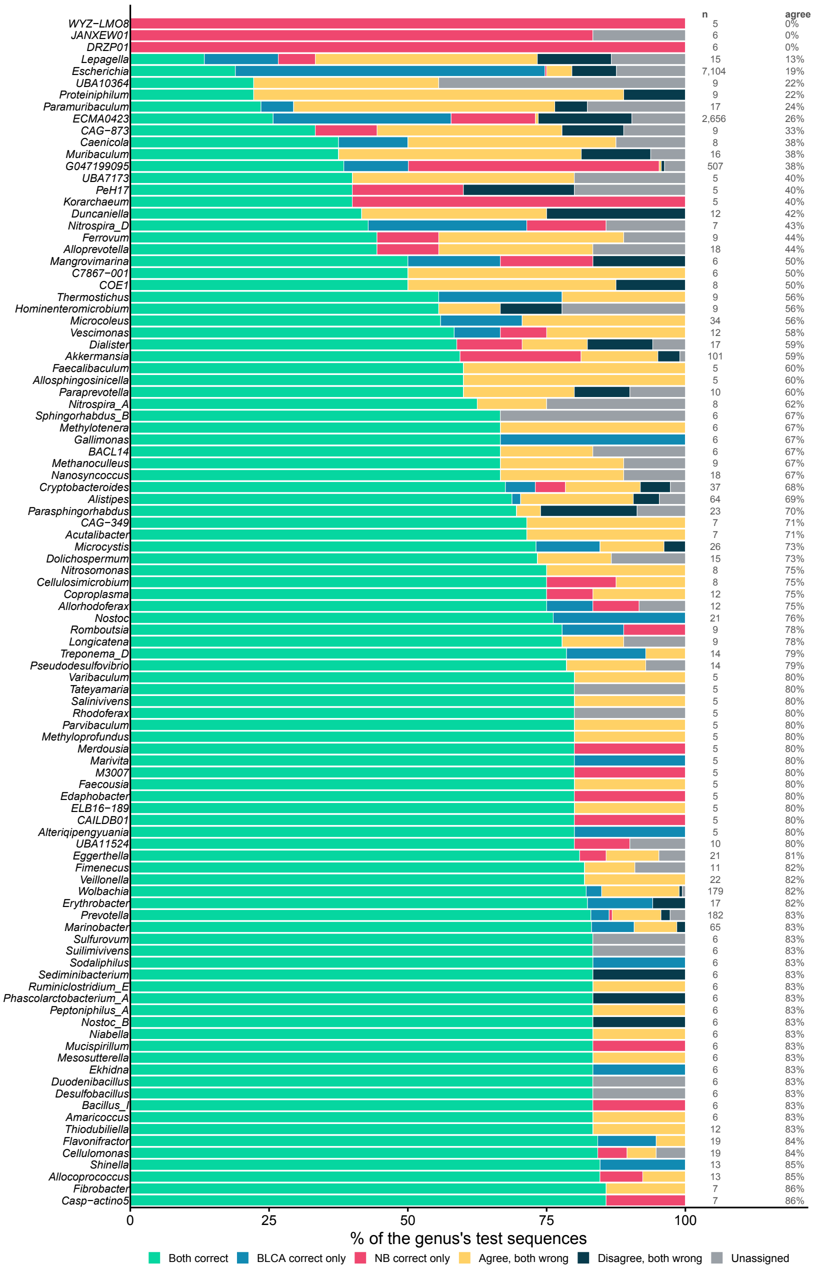


**Supplementary Figure S2. BLCA and naive Bayes genus calls compared, for sequences whose species is in the reference.** For each genus of the 16S rRNA gene GTDB r232 holdout, the percentage of that genus's test sequences on which the BLCA and naive Bayes genus calls were both correct, correct for BLCA only, correct for NB only, wrong in the same way (both classifiers returned the same wrong genus), wrong in different ways, or unassigned (at least one classifier made no assignment and neither was correct). Restricted to the 90,688 test sequences whose true species is present in the 90% training fraction. The hundred genera with the lowest agreement among those carrying at least five test sequences are shown, ordered by increasing agreement. n is the number of test sequences in the genus and agree the percentage on which both classifiers were correct. Every genus, including those below the five sequence floor, is listed in Supplementary Table S10.

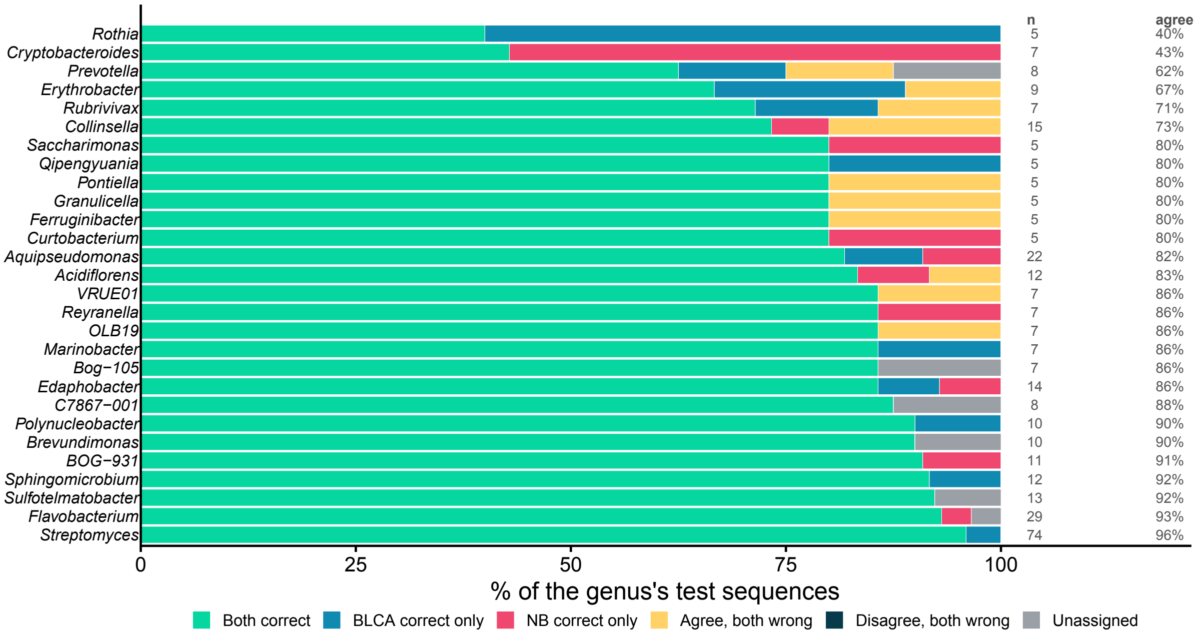


**Supplementary Figure S3. BLCA and naive Bayes genus calls compared, for sequences whose species is absent from the reference.** The same comparison as Supplementary Figure S2, restricted to the 5,043 test sequences whose true species is absent from the 90% training fraction. Of these, 4,130 have their genus present in the training fraction, so a correct genus call is possible. For the remaining 913 the genus is also absent, and they are excluded from the figure (771 of them, whose family is present, are scored at family level in Fig. 2c), though they are retained in Supplementary Table S11. Categories, ordering and the five sequence floor are as in Supplementary Figure S2.

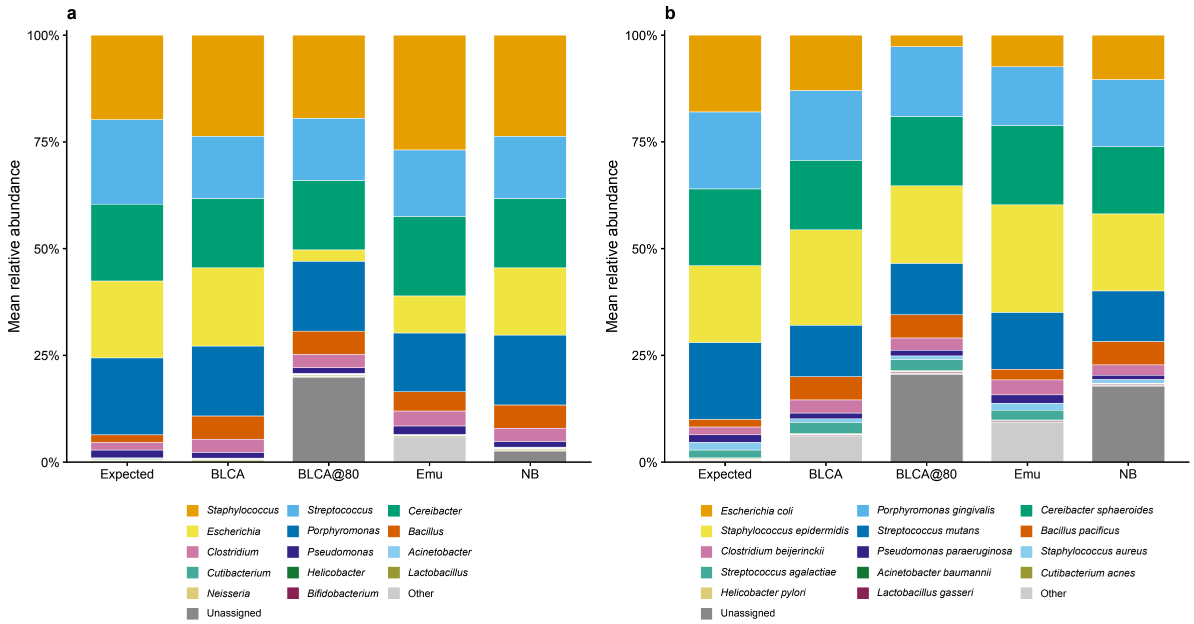


**Supplementary Figure S4. 16S rRNA gene ATCC MSA-1003 mock community composition.** Mean relative abundance across eight libraries at (a) genus and (b) species level, for the Expected (theoretical) composition and for each classifier (BLCA, BLCA@80, Emu, NB). Bars are stacked by taxon and read weighted, computed for Emu as the mean over all eight libraries of the per sample values in its genus and species tables (a taxon not reported in a library counts as 0) and for BLCA, BLCA@80 and NB by summing ASV read counts over all libraries at each rank and dividing by the total read count. Only the 14 most abundant expected taxa are coloured. The remaining expected taxa and every taxon absent from the mock are summed as Other, which therefore holds the *Mycobacterium* calls of Emu and the Escherichia boydii calls of BLCA (Supplementary Table S3 lists them). GTDB placeholder genera and named GTDB synonyms of mock members are folded into the corresponding mock organism. Emu is retained here because the ATCC mock carries known read abundances; the fungal ITS mock has no composition counterpart because it was pooled by equal volume and therefore has no abundance ground truth.

| | Expected | BLCA | BLCA@80 | Emu | NB |
|---|---|---|---|---|---|
| *Staphylococcus* | 19.8 | 23.7 | 19.5 | 26.9 | 23.7 |
| *Streptococcus* | 19.8 | 14.5 | 14.5 | 15.6 | 14.5 |
| *Cereibacter* | 18.0 | 16.3 | 16.3 | 18.6 | 16.3 |
| *Escherichia* | 18.0 | 18.4 | 2.7 | 8.7 | 15.8 |
| *Porphyromonas* | 18.0 | 16.3 | 16.3 | 13.7 | 16.3 |
| *Bacillus* | 1.8 | 5.5 | 5.5 | 4.5 | 5.5 |
| *Clostridium* | 1.8 | 3.1 | 3.1 | 3.5 | 3.1 |
| *Pseudomonas* | 1.8 | 1.3 | 1.3 | 2.0 | 1.3 |
| *Acinetobacter* | 0.2 | 0.2 | 0.2 | 0.2 | 0.2 |
| *Cutibacterium* | 0.2 | 0.1 | 0.1 | 0.0 | 0.1 |
| *Helicobacter* | 0.2 | 0.1 | 0.1 | 0.1 | 0.1 |
| *Lactobacillus* | 0.2 | 0.2 | 0.2 | 0.3 | 0.2 |
| *Neisseria* | 0.2 | 0.2 | 0.2 | 0.0 | 0.2 |
| *Bifidobacterium* | 0.0 | 0.0 | 0.0 | 0.0 | 0.0 |
| *Deinococcus* | 0.0 | 0.0 | 0.0 | 0.0 | 0.0 |
| *Enterococcus* | 0.0 | 0.0 | 0.0 | 0.0 | 0.0 |
| *Phocaeicola* | 0.0 | 0.0 | 0.0 | 0.0 | 0.0 |
| *Schaalia* | 0.0 | 0.0 | 0.0 | 0.0 | 0.0 |
| Unassigned | 0.0 | 0.0 | 19.9 | 0.0 | 2.6 |
| *Mycobacterium* | 0.0 | 0.0 | 0.0 | 5.0 | 0.0 |
| *Salmonella* | 0.0 | 0.0 | 0.0 | 0.3 | 0.0 |
| *Enterobacter* | 0.0 | 0.0 | 0.0 | 0.2 | 0.0 |
| *Klebsiella* | 0.0 | 0.0 | 0.0 | 0.1 | 0.0 |

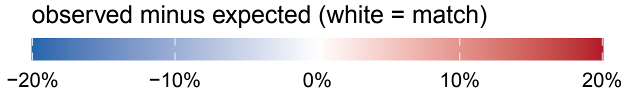


**Supplementary Figure S5. 16S rRNA gene ATCC MSA-1003 mock per genus relative abundance.** Heatmap of observed genus level relative abundance across classifiers (BLCA, BLCA@80, Emu, NB) and the Expected composition, averaged across eight libraries. Each cell value is the observed relative abundance of that genus, and cell colour encodes the deviation from the expected value (observed minus expected in percentage points) on a diverging scale, with white marking a match, warm colours an overcall and cool colours an undercall. The 18 mock genera are ordered by expected abundance. Below them, the Unassigned row gives the reads without a genus level call (19.9% for BLCA@80 and 2.6% for NB), and the last four rows are the genera absent from the mock (expected 0%) that a classifier reported at more than 0.1% of reads, all of them reported only by Emu. GTDB placeholder genera and named GTDB synonyms of mock members are folded into the corresponding mock organism.

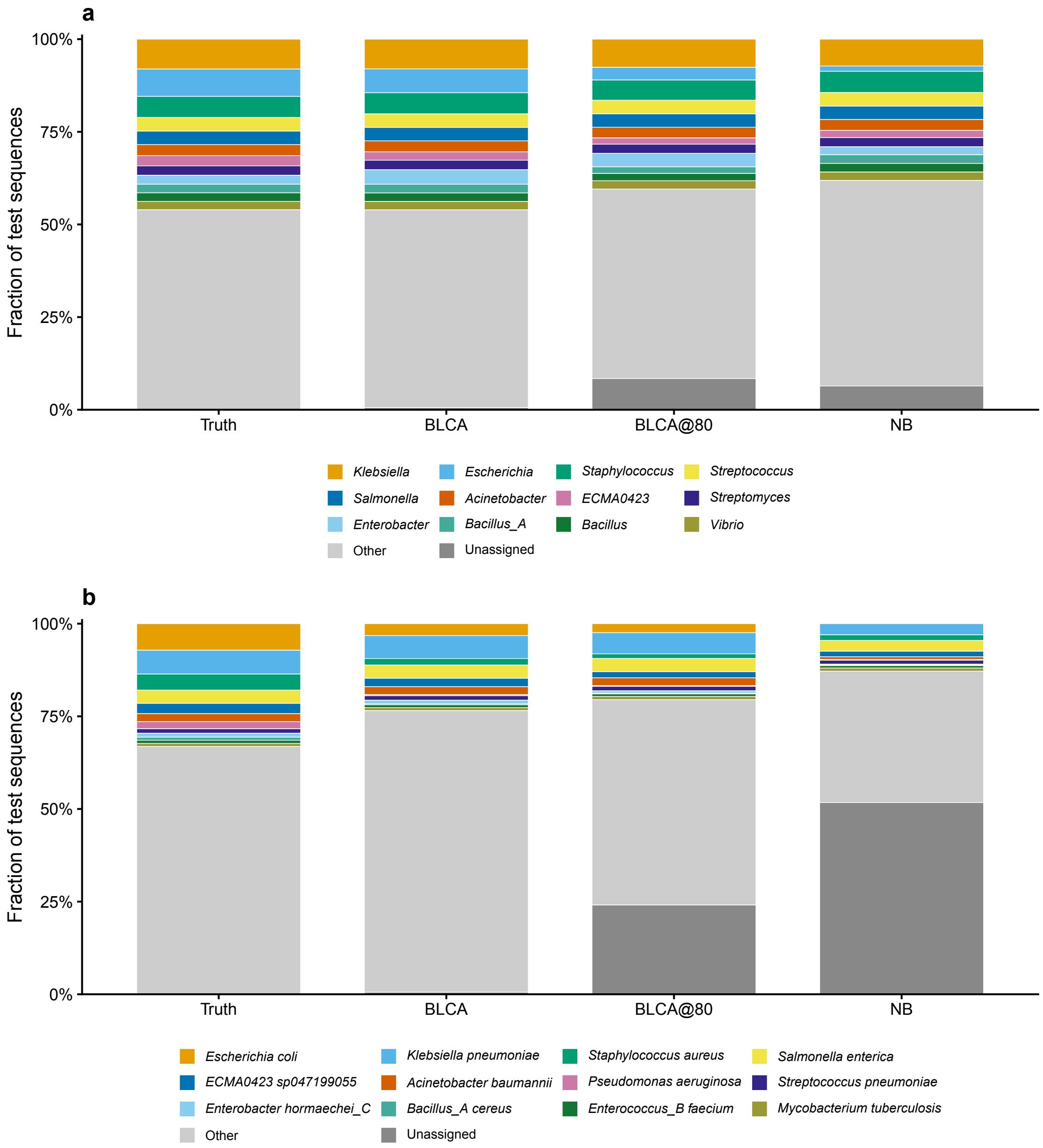


**Supplementary Figure S6. 16S rRNA gene holdout composition against the holdout truth.** Composition of the 16S rRNA gene GTDB r232 holdout test sequences at (a) genus and (b) species rank, comparing the holdout Truth (the GTDB annotation of each test sequence) against BLCA, BLCA@80 and the naive Bayes classifier (NB). Each test sequence counts once, so the bars give the fraction of test sequences assigned to each taxon. The top 12 taxa by truth abundance are shown, with the remainder lumped as Other and sequences receiving no call shown as Unassigned. Emu is absent because a holdout contains no reads to profile.

#### Supplementary References


1. Di Tommaso, P. et al. Nextflow enables reproducible computational workflows. Nature Biotechnology 35, 316–319 (2017). DOI: 10.1038/nbt.3820

2. Kurtzer, G. M., Sochat, V. & Bauer, M. W. Singularity: Scientific containers for mobility of compute. PLOS ONE 12, e0177459 (2017). DOI: 10.1371/journal.pone.0177459

3. Shen, W., Le, S., Li, Y. & Hu, F. SeqKit: A cross-platform and ultrafast toolkit for FASTA/Q file manipulation. PLOS ONE 11, e0163962 (2016). DOI: 10.1371/journal.pone.0163962

4. Martin, M. Cutadapt removes adapter sequences from high-throughput sequencing reads. EMBnet.journal 17, 10–12 (2011). DOI: 10.14806/ej.17.1.200

5. O'Brien, A., Lagos, C., Fernández, K., Ojeda, B. & Parada, P. ITSxRust: ITS region extraction with partial-chain recovery and structured diagnostics for long-read amplicon sequencing. Methods in Ecology and Evolution 17, 2843–2853 (2026). DOI: 10.1111/2041-210X.70393

6. Callahan, B. J. et al. DADA2: High-resolution sample inference from Illumina amplicon data. Nature Methods 13, 581–583 (2016). DOI: 10.1038/nmeth.3869

7. Bolyen, E. et al. Reproducible, interactive, scalable and extensible microbiome data science using QIIME 2. Nature Biotechnology 37, 852–857 (2019). DOI: 10.1038/s41587-019-0209-9

8. Curry, K. D. et al. Emu: Species-level microbial community profiling of full-length 16S rRNA Oxford Nanopore sequencing data. Nature Methods 19, 845–853 (2022). DOI: 10.1038/s41592-022-01520-4

9. Callahan, B. J., McMurdie, P. J. & Holmes, S. P. Exact sequence variants should replace operational taxonomic units in marker-gene data analysis. The ISME Journal 11, 2639–2643 (2017). DOI: 10.1038/ismej.2017.119

10. O'Brien, A., Lagos, C., Fernández, K., Ojeda, B. & Parada, P. EMITS: Expectation-maximization abundance estimation for fungal ITS communities from long-read sequencing. Methods in Ecology and Evolution (2026). DOI: 10.1111/2041-210X.70418

11. Parks, D. H. et al. GTDB release 10: A complete and systematic taxonomy for 715 230 bacterial and 17 245 archaeal genomes. Nucleic Acids Research 54, D743–D754 (2026). DOI: 10.1093/nar/gkaf1040

12. Parks, D. H. et al. GTDB: An ongoing census of bacterial and archaeal diversity through a phylogenetically consistent, rank normalized and complete genome-based taxonomy. Nucleic Acids Research 50, D785–D794 (2022). DOI: 10.1093/nar/gkab776

13. Abarenkov, K. et al. The UNITE database for molecular identification and taxonomic communication of fungi and other eukaryotes: Sequences, taxa and classifications reconsidered. Nucleic Acids Research 52, D791–D797 (2024). DOI: 10.1093/nar/gkad1039

14. Gao, X., Lin, H., Revanna, K. & Dong, Q. A Bayesian taxonomic classification method for 16S rRNA gene sequences with improved species-level accuracy. BMC Bioinformatics 18, 247 (2017). DOI: 10.1186/s12859-017-1670-4

15. Wang, Q., Garrity, G. M., Tiedje, J. M. & Cole, J. R. Naïve Bayesian classifier for rapid assignment of rRNA sequences into the new bacterial taxonomy. Applied and Environmental Microbiology 73, 5261–5267 (2007). DOI: 10.1128/aem.00062-07

16. Nouioui, I. et al. Genome-based taxonomic classification of the phylum Actinobacteria. Frontiers in Microbiology 9, 2007 (2018). DOI: 10.3389/fmicb.2018.02007

17. Tedersoo, L. & Anslan, S. Towards PacBio-based pan-eukaryote metabarcoding using full-length ITS sequences. Environmental Microbiology Reports 11, 659–668 (2019). DOI: 10.1111/1758-2229.12776

18. Tedersoo, L. et al. Benchmarking full-length ITS metabarcoding across Illumina 2 × 500, PacBio, and Oxford Nanopore sequencing using mock and soil communities. Molecular Ecology Resources 26, e70189 (2026). DOI: 10.1111/1755-0998.70189

19. Li, H. Minimap2: Pairwise alignment for nucleotide sequences. Bioinformatics 34, 3094–3100 (2018). DOI: 10.1093/bioinformatics/bty191

20. Liimatainen, K. et al. Taming the beast: A revised classification of Cortinariaceae based on genomic data. Fungal Diversity 112, 89–170 (2022). DOI: 10.1007/s13225-022-00499-9

21. McDonald, D. et al. Greengenes2 unifies microbial data in a single reference tree. Nature Biotechnology 42, 715–718 (2024). DOI: 10.1038/s41587-023-01845-1
