## Supplementary Figure 1 for "HiFiTaxa: a reproducible analysis pipeline for full length 16S rRNA gene and ITS amplicon sequencing from PacBio reads"

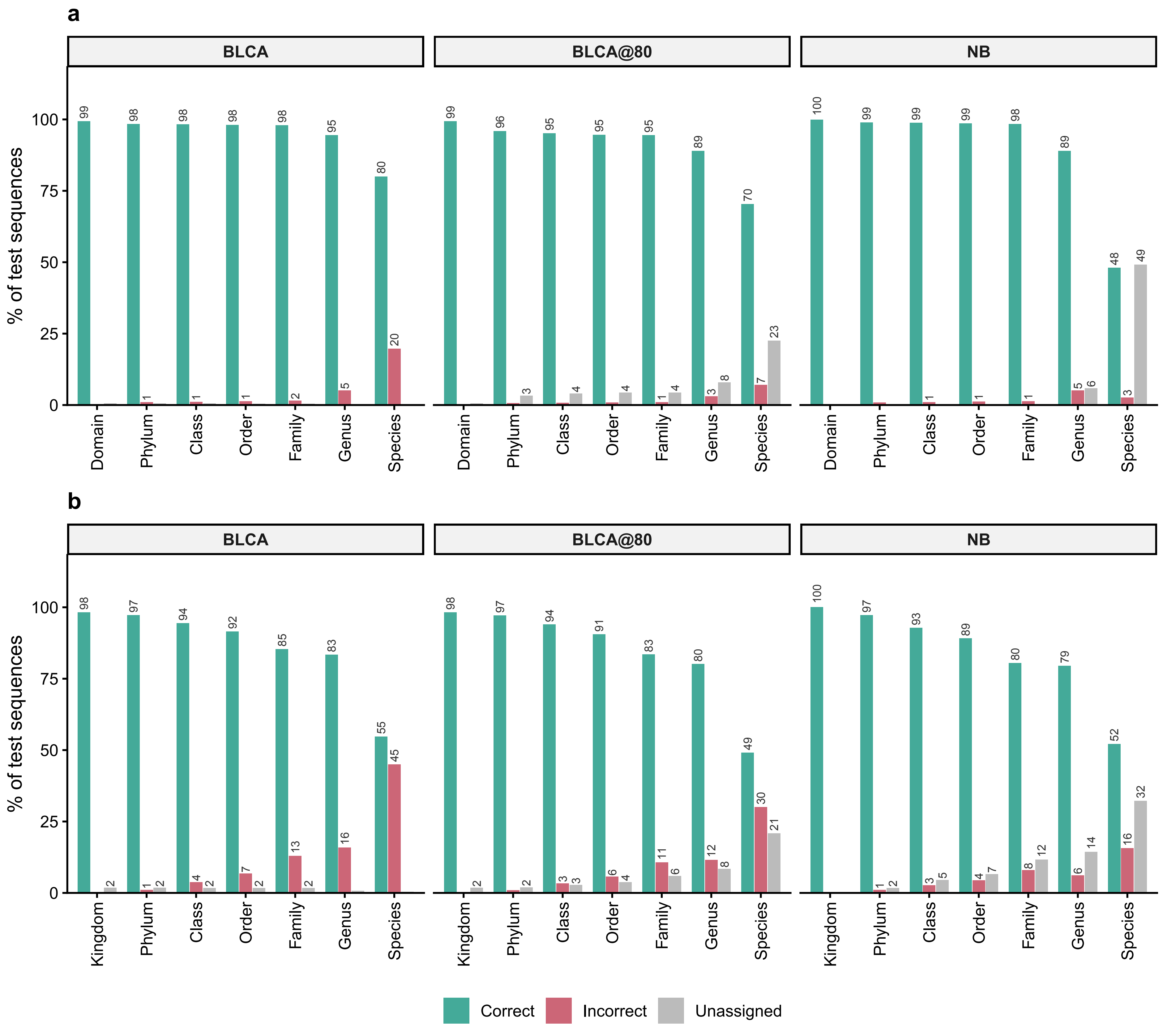
a
BLCA
BLCA@80
NB
% of test sequences
Domain
Phylum
Class
Order
Family
Genus
Species
b
Kingdom
Correct
Incorrect
Unassigned
