## Supplementary Figure 2 for "HiFiTaxa: a reproducible analysis pipeline for full length 16S rRNA gene and ITS amplicon sequencing from PacBio reads"

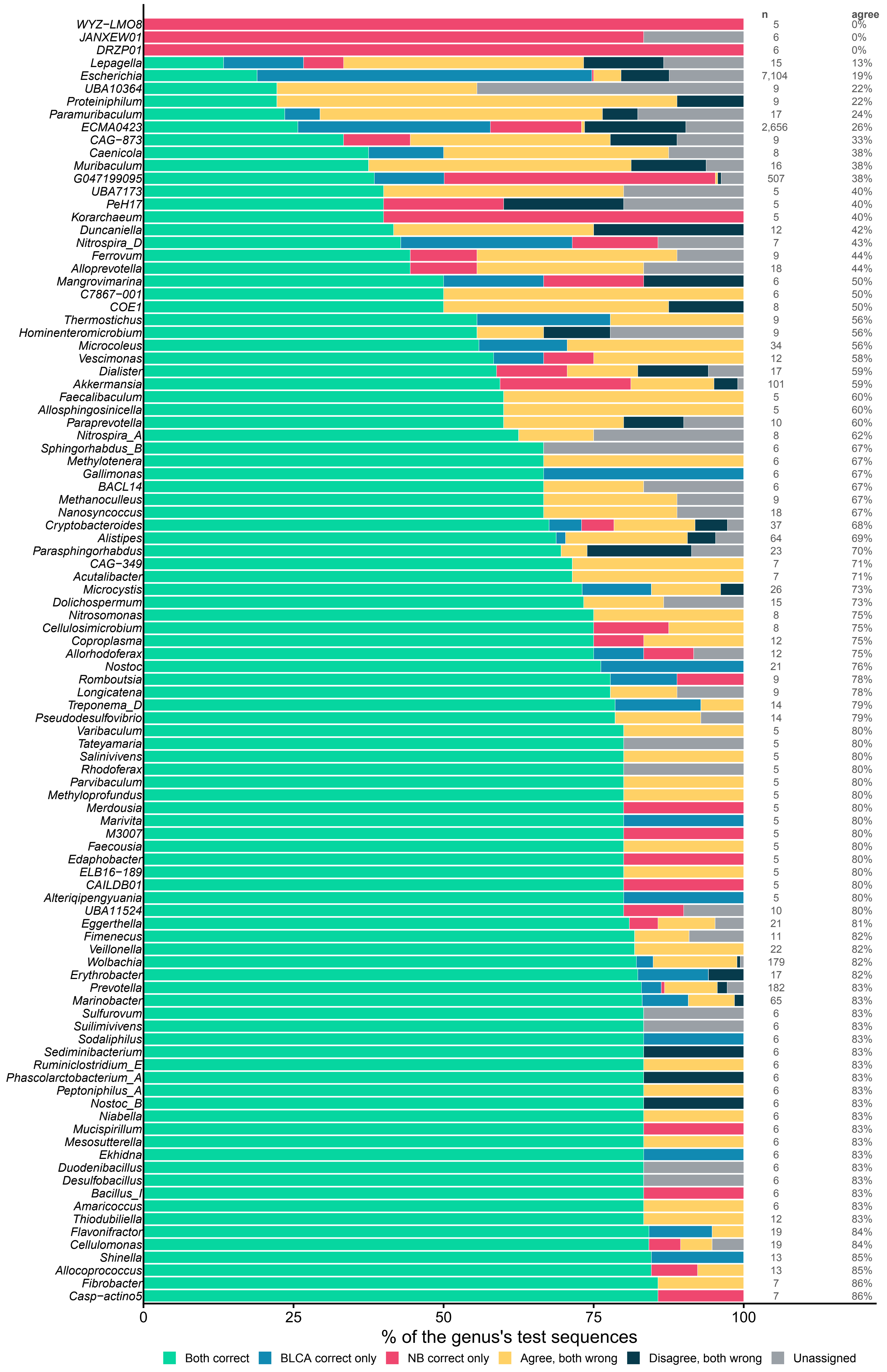

n
agree
WYZ−LMO8 5 0%
JANXEW01 6 0%
DRZP01 6 0%
Lepagella 15 13%
Escherichia 7,104 19%
UBA10364 9 22%
Proteiniphilum 9 22%
Paramuribaculum 17 24%
ECMA0423 2,656 26%
CAG−873 9 33%
Caenicola 8 38%
Muribaculum 16 38%
G047199095 507 38%
UBA7173 5 40%
PeH17 5 40%
Korarchaeum 5 40%
Duncaniella 12 42%
Nitrospira_D 7 43%
Ferrovum 9 44%
Alloprevotella 18 44%
Mangrovimarina 6 50%
C7867−001 6 50%
COE1 8 50%
Thermostichus 9 56%
Hominenteromicrobium 9 56%
Microcoleus 34 56%
Vescimonas 12 58%
Dialister 17 59%
Akkermansia 101 59%
Faecalibaculum 5 60%
Allosphingosinicella 5 60%
Paraprevotella 10 60%
Nitrospira_A 8 62%
Sphingorhabdus_B 6 67%
Methylotenera 6 67%
Gallimonas 6 67%
BACL14 6 67%
Methanoculleus 9 67%
Nanosyncoccus 18 67%
Cryptobacteroides 37 68%
Alistipes 64 69%
Parasphingorhabdus 23 70%
CAG−349 7 71%
Acutalibacter 7 71%
Microcystis 26 73%
Dolichospermum 15 73%
Nitrosomonas 8 75%
Cellulosimicrobium 8 75%
Coproplasma 12 75%
Allorhodoferax 12 75%
Nostoc 21 76%
Romboutsia 9 78%
Longicatena 9 78%
Treponema_D 14 79%
Pseudodesulfovibrio 14 79%
Varibaculum 5 80%
Tateyamaria 5 80%
Salinivivens 5 80%
Rhodoferax 5 80%
Parvibaculum 5 80%
Methyloprofundus 5 80%
Merdousia 5 80%
Marivita 5 80%
M3007 5 80%
Faecousia 5 80%
Edaphobacter 5 80%
ELB16−189 5 80%
CAILDB01 5 80%
Alteriqipengyuania 5 80%
UBA11524 10 80%
Eggerthella 21 81%
Fimenecus 11 82%
Veillonella 22 82%
Wolbachia 179 82%
Erythrobacter 17 82%
Prevotella 182 83%
Marinobacter 65 83%
Sulfurovum 6 83%
Suilimivivens 6 83%
Sodaliphilus 6 83%
Sediminibacterium 6 83%
Ruminiclostridium_E 6 83%
Phascolarctobacterium_A 6 83%
Peptoniphilus_A 6 83%
Nostoc_B 6 83%
Niabella 6 83%
Mucispirillum 6 83%
Mesosutterella 6 83%
Ekhidna 6 83%
Duodenibacillus 6 83%
Desulfobacillus 6 83%
Bacillus_I 6 83%
Amaricoccus 6 83%
Thiodubiliella 12 83%
Flavonifractor 19 84%
Cellulomonas 19 84%
Shinella 13 85%
Allocoprococcus 13 85%
Fibrobacter 7 86%
Casp−actino5 7 86%
0
25
50
75
100
% of the genus's test sequences
Both correct
BLCA correct only
NB correct only
Agree, both wrong
Disagree, both wrong
Unassigned
