## Supplementary Figure 3 for "HiFiTaxa: a reproducible analysis pipeline for full length 16S rRNA gene and ITS amplicon sequencing from PacBio reads"

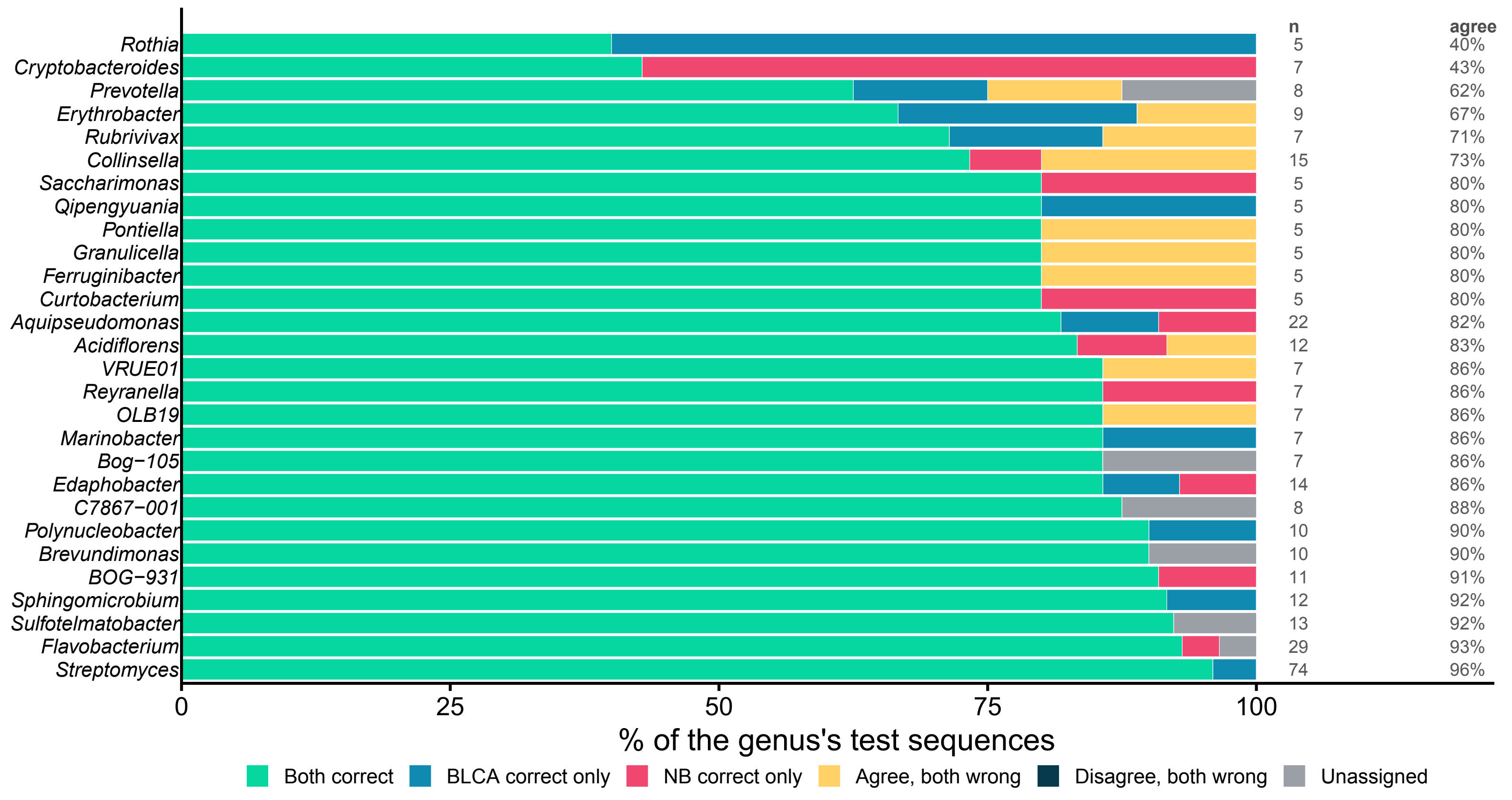

Rothia
Cryptobacteroides
Prevotella
Erythrobacter
Rubrivivax
Collinsella
Saccharimonas
Qipengyuania
Pontiella
Granulicella
Ferruginibacter
Curtobacterium
Aquipseudomonas
Acidiflorens
VRUE01
Reyranella
OLB19
Marinobacter
Bog−105
Edaphobacter
C7867−001
Polynucleobacter
Brevundimonas
BOG−931
Sphingomicrobium
Sulfotelmatobacter
Flavobacterium
Streptomyces
n
5
7
8
9
7
15
5
5
5
5
5
5
22
12
7
7
7
7
7
14
8
10
10
11
12
13
29
74
agree
40%
43%
62%
67%
71%
73%
80%
80%
80%
80%
80%
80%
82%
83%
86%
86%
86%
86%
86%
86%
88%
90%
90%
91%
92%
92%
93%
96%
0
25
50
75
100
% of the genus's test sequences
Both correct
BLCA correct only
NB correct only
Agree, both wrong
Disagree, both wrong
Unassigned
