## Supplementary Figure 4 for "HiFiTaxa: a reproducible analysis pipeline for full length 16S rRNA gene and ITS amplicon sequencing from PacBio reads"

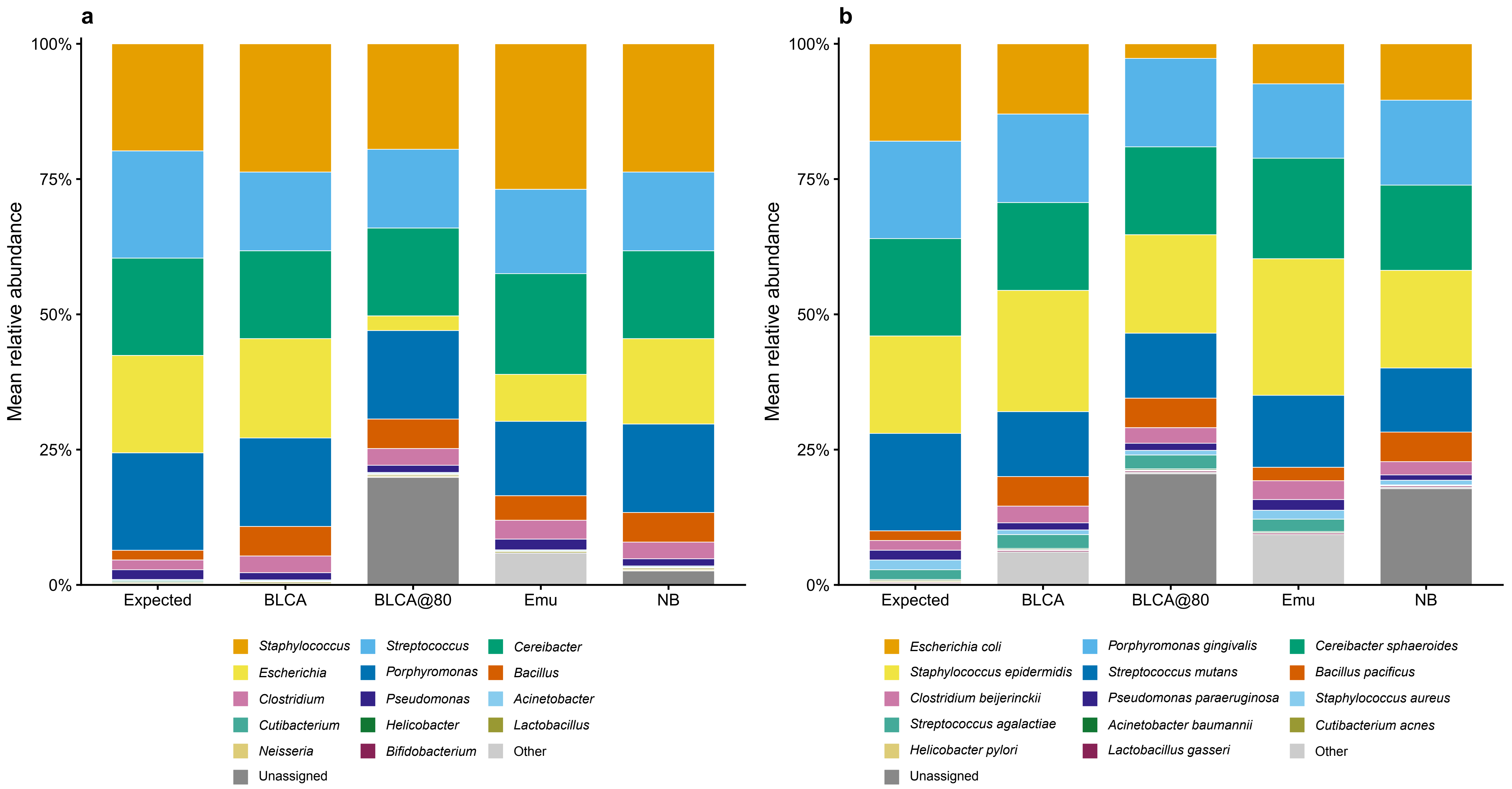
a
b
100%
75%
50%
25%
0%
Mean relative abundance
Expected
BLCA
BLCA@80
Emu
NB
Staphylococcus
Streptococcus
Cereibacter
Escherichia
Porphyromonas
Bacillus
Clostridium
Pseudomonas
Acinetobacter
Cutibacterium
Helicobacter
Lactobacillus
Neisseria
Bifidobacterium
Other
Unassigned
Escherichia coli
Porphyromonas gingivalis
Cereibacter sphaeroides
Staphylococcus epidermidis
Streptococcus mutans
Bacillus pacificus
Clostridium beijerinckii
Pseudomonas paraeruginosa
Staphylococcus aureus
Streptococcus agalactiae
Acinetobacter baumannii
Cutibacterium acnes
Helicobacter pylori
Lactobacillus gasseri
Other
Unassigned
