## Supplementary Figure 6 for "HiFiTaxa: a reproducible analysis pipeline for full length 16S rRNA gene and ITS amplicon sequencing from PacBio reads"

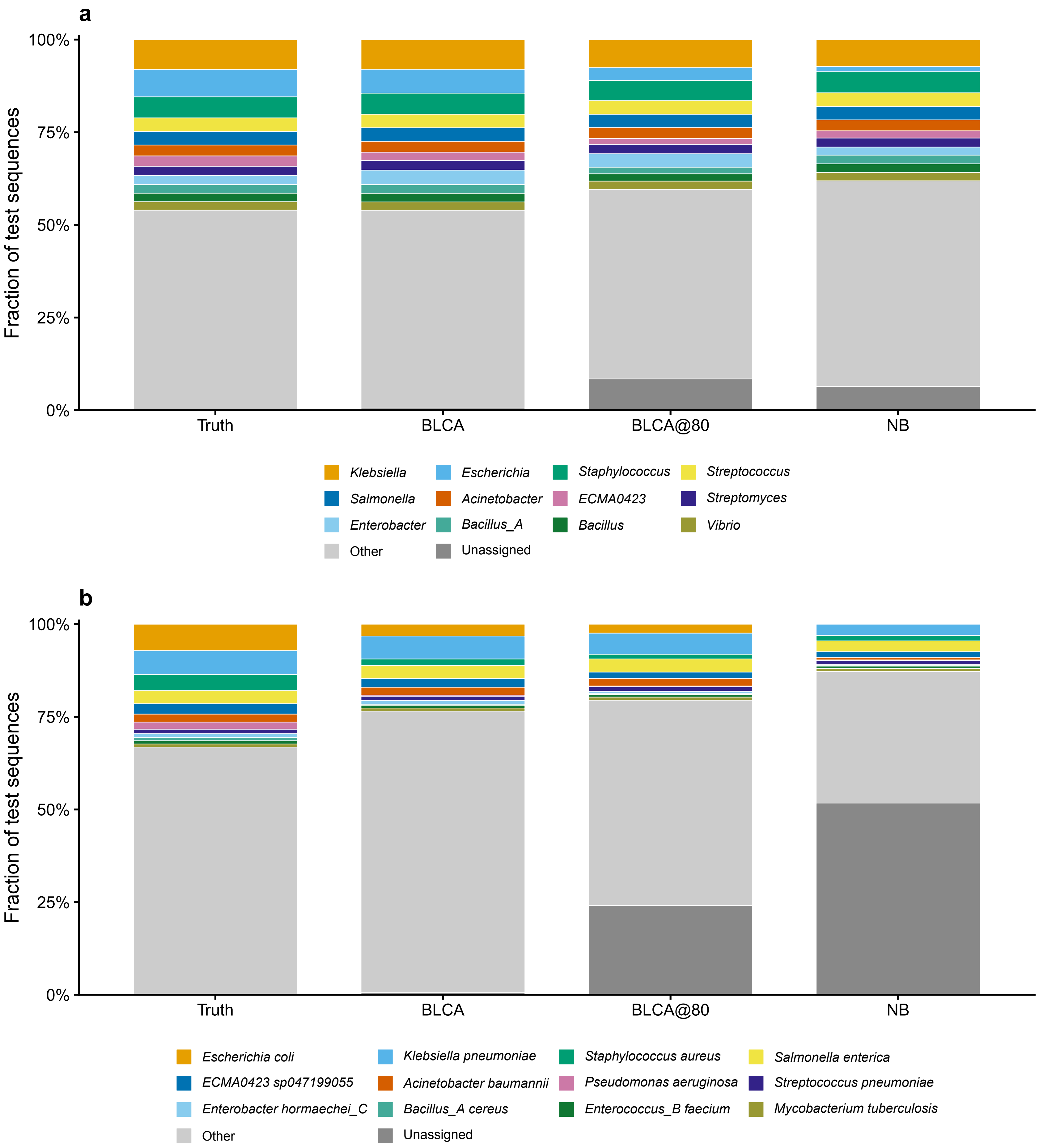
a
100%
75%
50%
25%
0%
Fraction of test sequences
Truth
BLCA
BLCA@80
NB
Klebsiella
Escherichia
Staphylococcus
Streptococcus
Salmonella
Acinetobacter
ECMA0423
Streptomyces
Enterobacter
Bacillus_A
Bacillus
Vibrio
Other
Unassigned
b
100%
75%
50%
25%
0%
Fraction of test sequences
Truth
BLCA
BLCA@80
NB
Escherichia coli
Klebsiella pneumoniae
Staphylococcus aureus
Salmonella enterica
ECMA0423 sp047199055
Acinetobacter baumannii
Pseudomonas aeruginosa
Streptococcus pneumoniae
Enterobacter hormaechei_C
Bacillus_A cereus
Enterococcus_B faecium
Mycobacterium tuberculosis
Other
Unassigned
